# Increases in therapist–patient physiological synchrony after dog–patient contact in canine-assisted psychotherapy: an exploratory pilot study

**DOI:** 10.64898/2026.10.06.756999

**Authors:** Stefanie Suttkus, Andy Schumann, Yubraj Gupta, Feliberto de la Cruz, Maria Geisler, Katrin Rieger, Steffen Schulz, Gerd Ganser, Karl-Jürgen Bär

## Abstract

**Introduction:** Canine-assisted psychotherapy (CAP) introduces a therapy dog into the therapist–patient relationship and may influence interpersonal processes underlying therapeutic change. This exploratory pilot study investigated whether direct patient–dog contact is associated with changes in autonomic arousal and therapist–patient physiological synchrony.

**Methods:** Skin conductance was recorded from 18 patients with depressive disorders and one therapist during 53 routine CAP sessions. A total of 192 episodes of direct patient–dog contact were identified from video recordings. Three-minute periods before and after each contact onsets were compared. Skin conductance level (SCL), skin conductance fluctuations (SCF), and therapist–patient physiological synchrony were assessed using zero-lag correlation and maximum cross-correlation. Sliding-window analyses examined temporal changes in synchrony, while randomly selected pseudo-events served as controls.

**Results:** SCL increased significantly following dog contact in both patients and the therapist, whereas SCF rate over 3 min did not change significantly. Therapist–patient physiological synchrony increased significantly after dog contact, as indicated by both zero-lag correlation and maximum cross-correlation. Sliding-window analyses confirmed these increases and showed a reduction in the temporal lag between the two physiological signals. Importantly, increases in synchrony were significantly greater following genuine dog-contact events than following pseudo-events.

**Discussion:** Direct physical contact between patients and therapy dogs is followed by increased autonomic arousal and enhanced therapist–patient physiological synchrony. These findings suggest that dog–patient contact may function as an interactional event that promotes physiological coordination and interpersonal attunement within the therapeutic dyad.

## INTRODUCTION

Psychotherapy is fundamentally an interpersonal process in which therapeutic change unfolds within the relational space shared by patient and therapist. Across diverse therapeutic orientations, the therapeutic alliance—comprising the affective bond between patient and therapist as well as their mutual agreement on therapeutic goals and tasks—is consistently identified as one of the most robust predictors of treatment outcome (Baier et al., 2020; Del Re et al., 2021; Flückiger et al., 2018). Establishing a trusting, secure, and collaborative relationship enables patients to engage meaningfully with demanding psychological processes, facilitates emotional processing, and provides a corrective interpersonal framework in which therapeutic transformation can occur (Rogers, 1957; Zilcha-Mano, 2017). However, despite widespread consensus regarding the clinical importance of the therapeutic alliance, the continuous, moment-to-moment interpersonal mechanisms through which this relationship develops, stabilizes, and shifts during sessions remain insufficiently understood.

A promising methodology for investigating these micro-process dynamics is the study of interpersonal synchronisation. Interpersonal synchronisation refers to the continuous temporal coordination of behavioural, physiological, or neural activity between interacting individuals (Hoehl et al., 2021). Conceptualised as a fundamental biological and behavioural substrate for social connectedness, empathy, and interpersonal co-regulation (Gordon and Bartsch, 2026; Mayo and Gordon, 2020), synchronisation in psychotherapy can manifest across multiple observational channels, including coordinated body movements, vocal prosody, and autonomic nervous system activity (Gordon et al., 2025; Kleinbub et al., 2020; Koole and Tschacher, 2016; Tourunen et al., 2022).

Physiological synchronisation is particularly valuable in clinical research because autonomic signals provide a continuous, implicit, and objective index of interpersonal regulation that cannot be fully captured via self-report measures or external behavioural coding (Gordon et al., 2025; Gordon and Bartsch, 2026; Mayo and Gordon, 2020). Empirical investigations have repeatedly linked therapist–patient physiological synchrony to central aspects of the therapeutic relationship, including alliance quality, in-session emotional depth, and overall treatment success (Bar-Kalifa et al., 2019; Gernert et al., 2024; Tschacher and Meier, 2020; Uhl et al., 2025)

Physiological synchrony is dynamic and may change in response to fluctuations in attention, affect, and arousal. Skin conductance, also referred to as electrodermal activity, provides a particularly sensitive index of sympathetic activation and can capture rapid, phasic changes in autonomic arousal (Boucsein et al., 2012; Dawson et al., 2007; Suttkus et al., 2021). This makes skin conductance relevant not only as a measure of physiological synchrony but also as a potential indicator of processes that may facilitate interpersonal coupling. Research suggests that arousal can shape attention and social information processing, while physiological arousal and interpersonal synchrony are closely related during social interaction (Critchley, 2002; Laine et al., 2009). For example, increased arousal has been associated with greater synchronisation of neural activity in attention-related networks, suggesting that arousal may influence the allocation of attention toward socially relevant information (Critchley, 2002; Zhang et al., 2014). Thus, transient changes in sympathetic arousal may represent a potential mechanism through which salient interpersonal events influence subsequent physiological coordination between interaction partners.

This contextual dependence is especially compelling when examining psychotherapy that incorporates animals. Canine-Assisted Psychotherapy (CAP) structurally expands the conventional dyadic therapist–patient relationship into a dynamic triadic interaction involving therapist, patient, and therapy dog. Therapy dogs are intentionally integrated into the therapeutic environment and have been shown to facilitate reductions in subjective and physiological stress while promoting positive emotional states (Faucher et al., 2025; Griffioen et al., 2023; Lundqvist et al., 2017). Clinical literature suggests that the presence of a therapy dog can enhance patient motivation, promote interpersonal trust, and accelerate therapeutic alliance formation (Hediger et al., 2021; Huber et al., 2024; Hüsgen et al., 2022; Lundqvist et al., 2017; Marciniuk et al., 2025; Pandey et al., 2024; Peel, 2025). Nevertheless, the precise process-level mechanisms through which the dog influences the therapist–patient dynamic remain largely unexamined.

The therapy dog may play a uniquely active role in interpersonal synchronisation because dogs possess sophisticated capacities for recognizing and responding to human social and emotional signals. Human–animal interaction studies demonstrate that dogs actively coordinate their attention and behaviour with human partners, exhibiting cross-species physiological and emotional coupling (Duranton et al., 2017, 2019; Katayama et al., 2019; Koskela et al., 2024; Lamontagne and Gaunet, 2024; Griffioen et al., 2020). Furthermore, direct tactile and social interaction with dogs induces immediate changes in human autonomic arousal and neurophysiological functioning (Beetz et al., 2012; Marti et al., 2022; Teo et al., 2024, 2022). Rather than serving merely as a passive ambient comfort, a therapy dog may operate as an active social catalyst that dynamically modifies the interactional ecology of the consulting room. Specifically, physical contact with the dog represents a salient interpersonal event capable of altering physiological arousal and restructuring subsequent autonomic coordination between patient and therapist.

This event-based dynamic is particularly relevant when dog–patient interaction becomes the focal point of subsequent therapeutic exploration. When the therapist and patient jointly observe, reflect upon, and discuss the patient’s interaction with the dog, the dog-related experience provides an immediate, experiential reference point (Ganser, 2017; Wohlfarth and Mutschler, 2026). Within this shared context, implicit emotional states, relational habits, and attachment patterns can be made explicit and explored safely. The physiological dynamics surrounding such a contact event can therefore illuminate how the primary therapeutic relationship responds to and integrates an experience involving the third member of the therapeutic triad.

From a dynamical systems perspective, dog–patient contact can be conceptualized as a discrete interactional perturbation, a potential transition point within the therapeutic encounter. Contact with the therapy dog may acutely alter the autonomic baseline of one or both participants, subsequently reorganizing patterns of therapist–patient physiological coupling. Measuring skin conductance synchrony before and after dog–patient contact allows researchers to evaluate whether localized triadic events induce systemic shifts in dyadic autonomic coordination. Rather than treating synchrony merely as a static, session-averaged trait, an event-related design provides a temporal window into process-level relational shifts.

The present exploratory pilot study addresses this gap by evaluating whether moments of dog–patient contact are associated with an increased patient sympathetic arousal and elevated therapist–patient physiological synchrony. By conducting a comparative, event-centered analysis of pre- and post-contact autonomic dynamics, this study offers an initial process-oriented investigation into how interaction with a therapy dog influences real-time physiological coordination within the therapeutic alliance.

## METHODS

Skin conductance recordings were collected during routine CAP sessions conducted in a clinical setting. The therapy dogs were two male Large Munsterlander aged 5 years (24 sessions) and 11 years (14 sessions). Sessions involved one therapist and varying patients and were performed without experimental manipulation or behavioral constraints, allowing spontaneous variation in movement, posture, and interaction behavior representative of routine therapeutic practice.

Eighteen patients (15 females, 3 males, 35 ± 10 years of age, range 22-55 years) were included that have been diagnosed with a depressive disorder and different comorbidities. In total, 53 sessions were analyzed involving these patients with each session lasted 45 to 70 minutes. All sessions were conducted by a licensed psychological psychotherapist with formal training and experience in dog-assisted psychotherapy. The therapeutic procedures followed the clinic’s routine practice and were not modified for the purposes of physiological recording. Prior to participation, all patients provided written informed consent for participation in the study and for audiovisual and physiological recordings. The study protocol was approved by the Ethics Committee of the Medical Faculty of Friedrich-Schiller University Jena (2025-3882_1-BO-A).

### Acquisition and preprocessing of skin conductance

Skin conductance was recorded from patients and the therapist using wireless devices attached to the forearm (PPGED-R, BioNomadix, BIOPAC Systems Inc., Goleta, CA, USA). Two gelled Ag/AgCl electrodes were attached to the balm of the left hand at the proximal and distal phalanx. Signals were digitalized simultaneously by multi-channel amplifier system at 1000 Hz (MP160, BIOPAC Systems Inc., Goleta, CA, USA)

From the raw signals we estimated skin conductance level and the number of skin conductance fluctuations (Schumann 2017). To estimate physiological synchrony, we extracted the phasic components of skin conductance traces. First, raw signals were downsampled to 40 Hz for core preprocessing (including anti-aliasing filter). The tonic signal component was isolated using a 4th-order low-pass Butterworth filter with a cutoff frequency of 5 Hz, and the phasic component was derived via subtraction of the tonic part from the downsampled signal. An additional 4th-order low-pass filter (f_c_ = 0.5Hz) was applied to smooth the phasic signal. Finally, the smoothed phasic component was downsampled to 5 Hz for synchrony calculation and standardized using within-session z-score normalization.

Recordings with low signal quality were dropped after visual inspection. Segments with transient loss of the signal were excluded from the analysis.

### Video recordings and detected events

All sessions were video recorded using two synchronized digital cameras (50fps, HDR- CX240, Sony Corporation, Tokyo, Japan) positioned on opposite sides of the therapy room to ensure continuous visual coverage of both dogs throughout each session Synchronization between video streams and physiological recordings was achieved using an LED light which automatically turned on when the physiological recording was started and could be identified on video.

We coded situations in which direct physical contact occurred between the patient and the therapy dog retrospectively from video recordings. One investigator (SS) performed the primary annotation, which were independently reviewed by a second investigator (AS). Disagreements (deviation by more than 30s) were resolved by consensus. The onset of these physical contacts were coded as events and temporally aligned with the physiological recordings.

For each event, a continuous segment surrounding the event was extracted, consisting of 180 s before and 180 s after event onset. Thus, each event was represented by a 6-min analysis period extending from ™180 to +180 s relative to the onset. The *Pre* event period was defined as the 180-s interval preceding event onset, and the *Post* event period as the 180-s interval following event onset. Whenever windows around events included artifacts or quality exclusions, these events were excluded from further analysis.

### Whole-segment synchrony

Interpersonal Skin conductance synchrony was quantified separately for the *Pre* and *Post* event periods. Two complementary correlation-based measures were calculated. First, zero-lag Pearson correlation was calculated between the patient and therapist skin conductance signals.

Second, cross-correlation analysis was performed to quantify synchrony occurring at temporal lags. Cross-correlation was calculated over a maximum lag of ±5 s. The maximum positive cross-correlation was extracted together with the corresponding temporal lag. Thus, the maximum cross-correlation quantified the strongest positive patient–therapist correspondence within the specified lag range, whereas the optimal lag indicated the temporal offset at which this maximum occurred.

### Sliding-window synchrony analysis

To examine the temporal evolution of interpersonal synchrony around behavioral events, a sliding-window analysis was performed on the continuous 360-s event-centered segment (Boker et al., 2002). Windows of 30 s were advanced in 5-s increments, yielding overlapping windows spanning the pre- and post-event periods (Prochazkova et al., 2022).

For each valid window, zero-lag correlation, maximum positive cross-correlation, and the lag corresponding to the maximum cross-correlation were calculated using the same procedures as for the whole-segment analysis. Sliding windows were assigned to the pre-event period when their center occurred before event onset and to the post-event period when their center occurred at or after event onset.

For each event, window-level measures were averaged separately across all *Pre* and *Post* event windows. These event-level averages were subsequently used to quantify changes in synchrony from before to after the event.

### Random pseudo-event analysis

To confirm that physiological changes around coded events were not unspecific artifacts, real event windows were compared against pseudo-event control windows sampled across non-event time points (Schumann et al., 2020; Steinfath et al., 2025). Equivalent non-event control windows (3 min) were sampled across sessions to establish a comparative baseline (Martin and Stelmaczonek, 1988).

Pseudo-event onset times were selected randomly within each recording session rather than via a systematic temporal pattern or behavioral matching. Up to 20 pseudo-events were generated per session and fewer were accepted when eligible recording time was limited. To prevent contamination, *Pre* and *Post* event segments of pseudo-events were prohibited from overlapping with one another or with real events.

Pseudo-events were processed through the exact same analysis pipeline as genuine behavioral events, calculating 180-s pre- and post-event whole-segment and sliding-window synchrony measures, event-level sliding-window averages, and pre–post change scores.

### Statistical analysis

Raw skin conductance was analyzed around events using 3-min pre- and post-event windows. Primary outcomes were skin conductance level and the number of skin conductance fluctuations (Schumann et al., 2017). Linear mixed-effects models tested pre-to-post changes separately for patients and therapists, accounting for repeated observations within patients and sessions.

Physiological synchrony was quantified using changes in correlation-based measures between *Pre*- and *Post*-event periods. For genuine events, *Pre*-to-*Post* changes were assessed using linear mixed-effects models with maximum cross-correlation as the primary outcome. Because change scores were defined as post-event minus pre-event values, the model intercept represented the mean change. Patient-specific random intercepts were included, with session-level variance components nested within patient. Models were estimated using restricted maximum likelihood (REML).

To assess event-specific synchrony beyond temporal coincidence, genuine events were compared with pseudo-events using mixed-effects models including event class (genuine vs. pseudo) as a fixed effect, with pseudo-events as the reference category and the same patient/session random-effects structure. The coefficient for event class represented the estimated difference in synchrony change between genuine and pseudo-events.

## RESULTS

### Basic characteristics of detected events

Across all 53 sessions, a total number of 192 events have been detected coding a physical contact between dog and patient. On average 3.6 ± 2.7 events per session were detected. The duration was 1.06 ± 1.48 min on average (see **Figure 1**).

**Figure 1.**
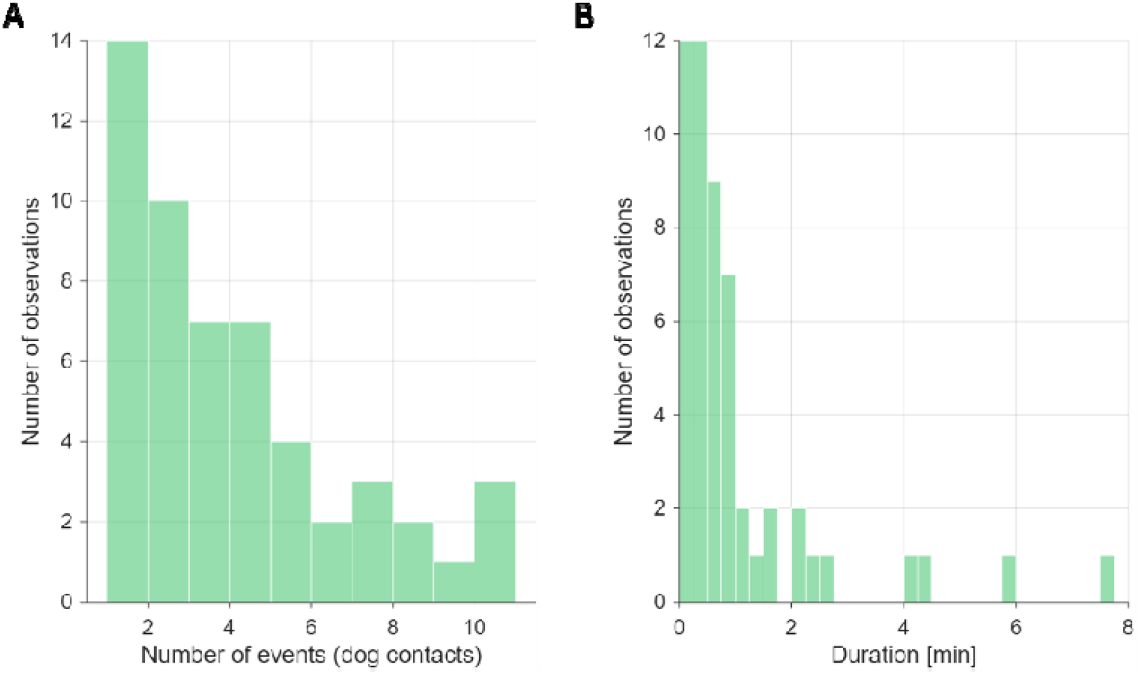
Descriptive statistics of detected events coding dog-patient contact. A: Number of coded events per session. B: Duration of the contact in minutes.

### Changes in skin conductance in both patients and therapist after events

There was a significant effect of event phase (*Pre*/*Post*) on SCL in both partners (see **Figure 2**). SCL increased significantly from *Pre* to *Post* in patients (β = 0.927, 95% CI [0.080, 1.773], *p* = 0.032) as well as in the therapist (β = 0.588, 95% CI [0.120, 1.057], *p* = 0.014). In contrast, event phase did not show a statistically significant effect on the number of fluctuations (SCF) in either patients (β = 0.443, 95% CI [™0.471, 1.356], *p* = 0.342) or the therapist (β = ™0.219, 95% CI [™1.057, 0.620], *p* = 0.609).

**Figure 2.**
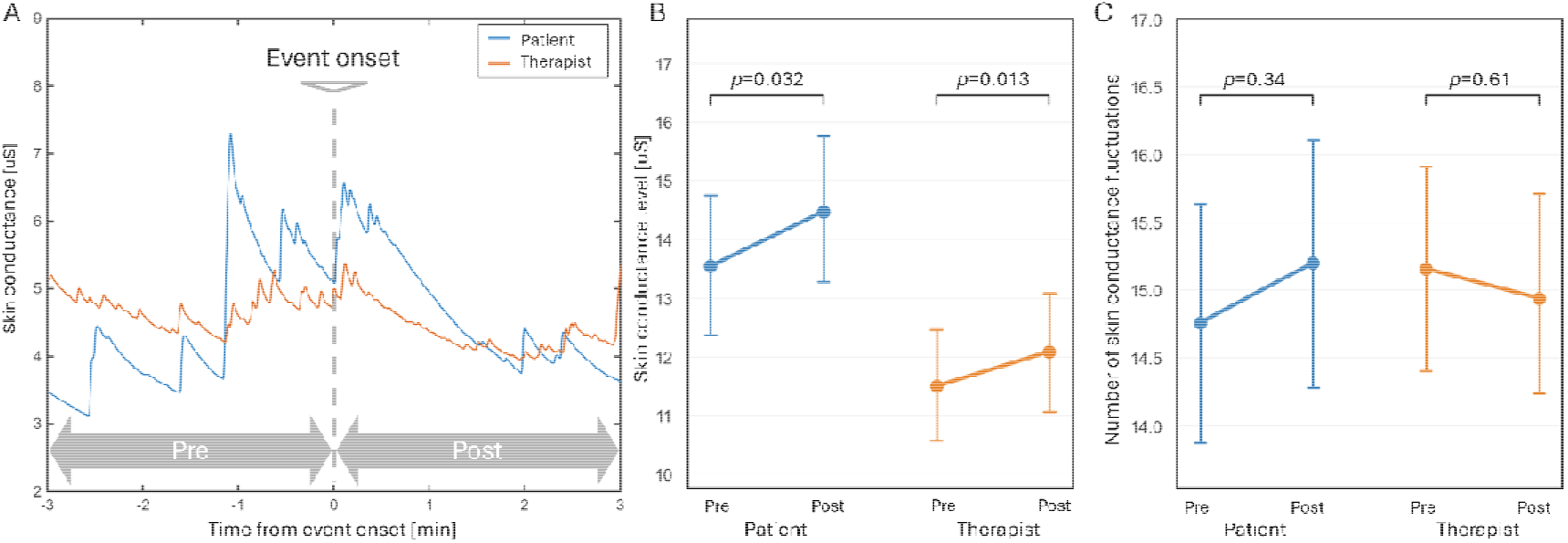
Changes in individual skin conductance after events. A: Example of a skin conductance recording aligned with an event (onset of direct contact between patient and dog). *Pre* and *Post* segments of three minutes each were analyzed. B: Skin conductance levels increased from before (*Pre*) to after the event (*Post*) in patients (blue) and the therapist (orange). C: Skin conductance fluctuations did not change significantly from before (*Pre*) to after the event (*Post*) in either patients (blue) or the therapist (orange). Error bars depict 95% CI of the estimated marginal means of linear mixed models.

### Skin conductance synchrony after direct dog-contact

To check for Pre–Post changes of physiological synchrony we set up linear mixed-effects models for correlation measures of skin conductance between patients and the therapist over the whole *Pre* and *Post* segments lasting 3 minutes. We discovered a significant increase in bivariate zero-lag correlation (β = 0.102, 95% CI [0.038, 0.166], *p* = 0.002) as well as the maximum cross correlation (β = 0.093, 95% CI [0.048, 0.139], *p* < 0.001). In contrast, the *Pre*–*Post* change in the optimal lag was not statistically significant (β = ™0.555, 95% CI [™1.385, 0.274], *p* = 0.190).

When these measures of synchrony were estimated in moving time windows, we saw a similar pattern (**Figure 3**). Averaged zero-lag correlation (β = 0.088, 95% CI [0.062, 0.114], *p* < 0.001) and the maximum cross-correlation showed a significant increase form *Pre* to *Post* event phase (β = 0.111, 95% CI [0.073, 0.149], *p* < 0.001). The optimal lag decreased significantly from *Pre* to *Post* event (β = -0.240, 95% CI [-0.484, -0.005], *p* = 0.036).

**Figure 3.**
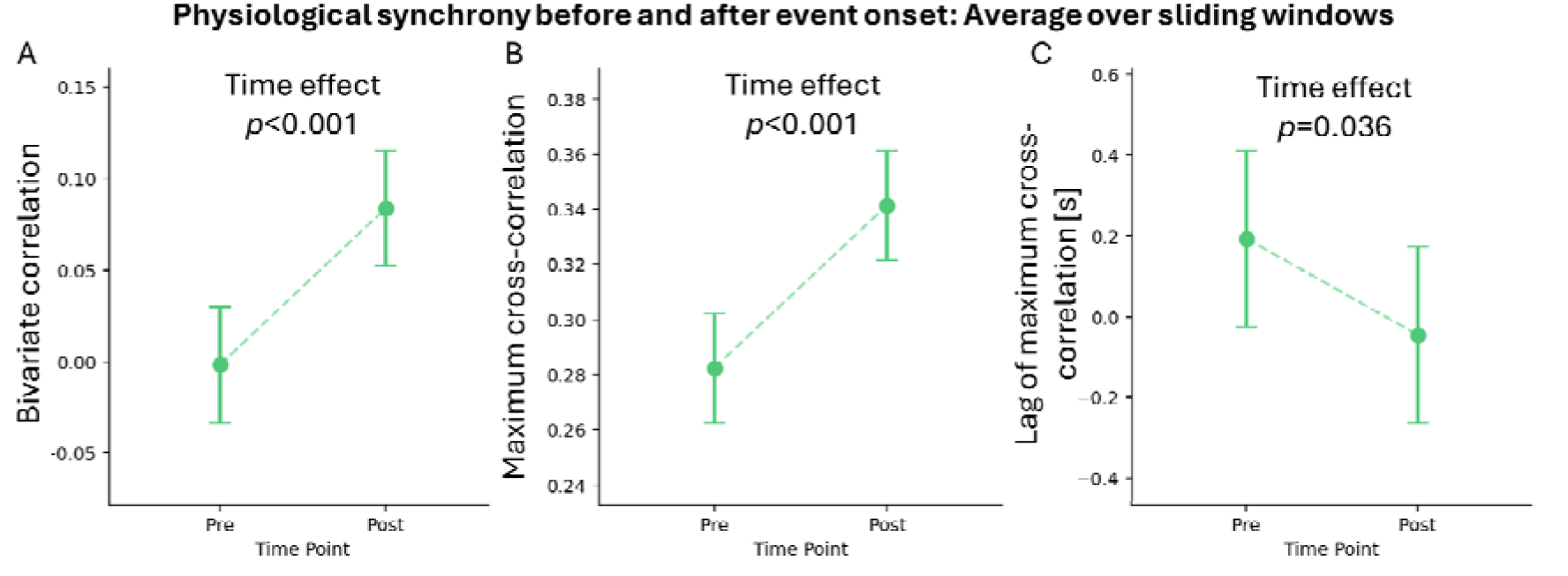
Physiological synchrony measured by sliding window cross-correlation of skin conductance before (*Pre*) and after (*Post*) an event. A: Average correlation with zero lag increased from *Pre* to *Post* event. B: Averaged maximum cross-correlation also increased form *Pre* to *Post* event. C: The average lag at with correlation was highest decreased from *Pre* to *Post*. Error bars depict 95% CI of the estimated marginal means of linear mixed models.

### Specificity analysis using pseudo-events

To investigate the specificity of the event based changes in physiological synchrony, we defined pseudo-events randomly throughout the sessions where no real events occurred. A comparison of *Pre*-*Post* changes between real and pseudo-events showed a significantly greater increase in zero-lag correlation following real events compared with pseudo-events (β = 0.080, 95% CI [0.006, 0.153], *p* = 0.033). Likewise, maximum cross-correlation exhibited a significantly greater *Pre–Post* change for real events than for pseudo-events (β = 0.087, 95% CI [0.017, 0.156], *p* = 0.014). In contrast, the difference between real and pseudo-events in the *Pre–Post* change in the optimal lag was not statistically significant (β = -0.841, 95% CI [™1.991, 0.309], *p* = 0.152).

When looking at sliding window correlation results, we found significantly greater increases in synchrony measures after real events (see **Figure 4**). Specifically, the *Pre–Post* change in zero-lag correlation (β = 0.087, 95% CI [0.043, 0.132], *p* < 0.001), as well as the maximum cross-correlation were significantly greater for real events than for pseudo-events (β = 0.062, 95% CI [0.034, 0.091], *p* < 0.001). Finally, the change of the optimal lag did not differ between real events and pseudo-events (β = -0.180, 95% CI [-0.568, 0.208], *p* = 0.363).

**Figure 4.**
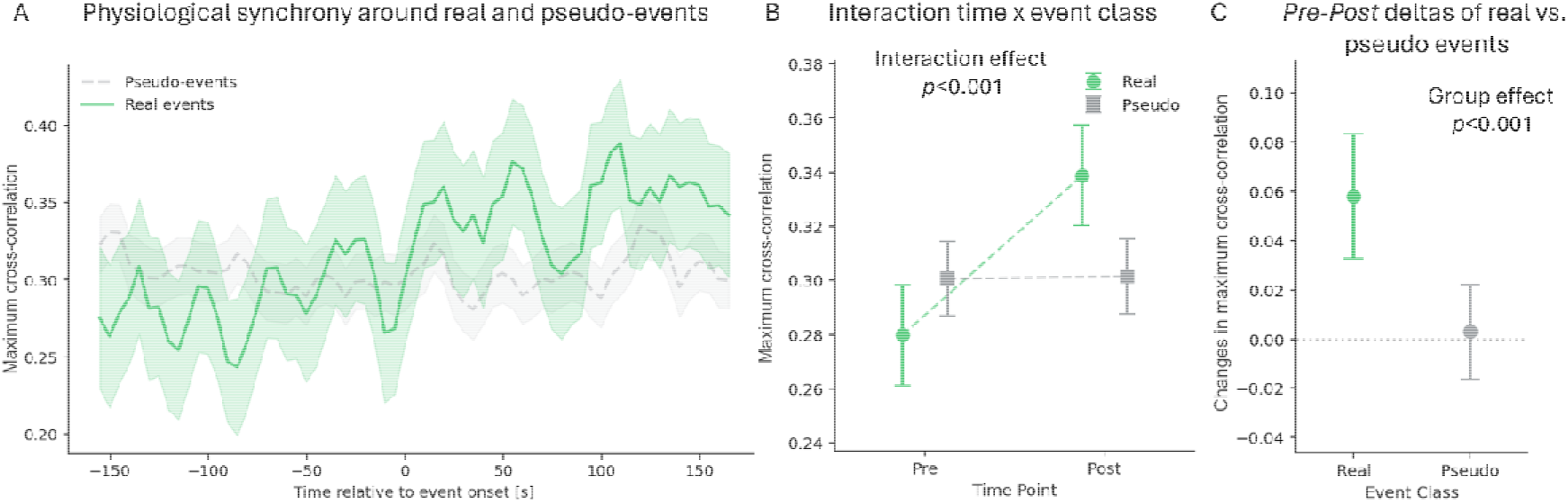
Comparison of physiological synchrony around real events (green) when compared to pseudo-events (gray). A: Time course of averaged maximum cross-correlation before and after real events and pseudo-events (at t=0). B: Sliding window maximum cross-correlation in the *Pre* and *Post* phases of real events and pseudo-events. C: Comparison of the *Pre-Post* changes in sliding window maximum cross-correlation around real events and pseudo-events. Error bars depict 95% CI of the estimated marginal means of linear mixed models.

In **Figure 4A**, the average time-resolved synchrony index before and after the event is depicted. Although synchrony fluctuates considerably over time, an elevation of synchrony after the onset of the event (at t=0) is clearly visible. Synchrony seems to remain elevated over the analysis window of 3 min.

## DISCUSSION

The primary objective of this exploratory pilot study was to examine whether direct physical contact between a therapy dog and a patient during Canine-Assisted Psychotherapy (CAP) alters therapist–patient autonomic dynamics. Specifically, we investigated whether dog– patient contact functions as an interactional transition point that enhances therapist–patient skin conductance synchrony. Our findings demonstrate that following direct dog–patient contact: (1) individual skin conductance levels (SCL) increased significantly in both patients and the therapist, while skin conductance fluctuations (SCF) remained unchanged; (2) therapist–patient physiological synchrony, measured via bivariate zero-lag correlation and maximum cross-correlation, increased significantly; and (3) these post-event increases in synchrony were specific to actual dog-contact events compared to randomly assigned pseudo-events. Together, these results provide initial empirical evidence that physical interaction with a therapy dog actively restructures the real-time autonomic coordination of the therapeutic dyad.

### Autonomic Arousal Following Dog–Patient Contact

The observed increase in skin conductance levels (SCL) in both patient and therapist following dog contact suggests a heightened state of sympathetic activation. Electrodermal activity, particularly SCL, is a reliable index of sympathetic nervous system arousal associated with emotional processing, cognitive engagement, and attentional allocation (Gernert et al., 2024; Kleinbub et al., 2020). The increase in patient SCL aligns with prior human–animal interaction research indicating that physical contact with a dog induces acute physiological and neural activation (Marti et al., 2022; Teo et al., 2024). Rather than acting purely as a tranquilizing agent that reduces arousal, contact with a therapy dog appears to represent a salient, engaging event that elevates physiological readiness and affective focus within the session.

Interestingly, the therapist also exhibited a significant increase in SCL following patient–dog contact. Within the triadic framework of CAP, the therapist is not an isolated observer; rather, the dog’s interaction with the patient provides a prominent behavioral and emotional signal within the scene. The simultaneous elevation in therapist SCL likely reflects increased attentional tracking, heightened emotional attunement to the patient’s state, or shared co- arousal as the dyad shifts its collective focus to the interaction involving the dog.

Notably, skin conductance fluctuations, which capture phasic spontaneous non-specific or transient responses, did not change significantly for either participant. As physiological synchrony was estimated on the phasic component of skin conductance, the elevated tonic level is not driving mechanistically the increases in synchrony. This selective increase in tonic SCL alongside stable phasic activation may suggest a state of sustained, organized physiological engagement rather than acute distress or erratic arousal.

### Enhancement of Therapist–Patient Physiological Synchrony

Beyond individual physiological changes, our findings reveal a significant increase in therapist–patient skin conductance synchrony following dog–patient contact. Across both full-segment models (3-minute windows) and time-resolved sliding window analyses, zero-lag correlation and maximum cross-correlation increased significantly post-event. In sliding window models, the optimal lag between patient and therapist signals also decreased, indicating tighter temporal coupling in sympathetic activity.

In dyadic psychotherapy, physiological synchrony has been conceptualized as an implicit biological substrate for social attunement, empathy, and emotional co-regulation (Gordon and Bartsch, 2026; Hoehl et al., 2021; Palumbo et al., 2017). Previous research has linked higher therapist–patient autonomic coupling to stronger therapeutic alliance ratings and deeper emotional engagement (Bar-Kalifa et al., 2019; Koole and Tschacher, 2016; Tschacher and Meier, 2020; Uhl et al., 2025). The specificity analysis using pseudo-events further reinforces these findings: real dog-contact events yielded significantly greater increases in both lag-zero and maximum cross-correlation compared to control baseline periods. This confirms that the observed enhancement of physiological coupling was not an artifact of time-on-task or general session progression, but was specifically driven by the occurrence of dog–patient contact.

From a dynamical systems perspective, the therapy dog appears to act as a social catalyst and a perturbation event that shifts the dyadic interactional state. In conventional human-only psychotherapy, interpersonal synchrony fluctuates dynamically based on conversational topics, nonverbal attunement, and affective intensity (Tourunen et al., 2022). Introducing a third, non-human social partner alters this dynamic, turning a dyadic system into a fluid triad (Mittly et al., 2024). Contact with the dog provides a concrete, shared experiential anchor for both patient and therapist. As both participants jointly perceive, process, and discuss the interaction with the dog, their autonomic nervous systems appear to align more closely in time. The dog-related experience may thus function as a bridge, facilitating shared attention and emotional co-regulation between patient and therapist (Ganser, 2017; Hediger et al., 2021).

### Limitations and Future Directions

Several limitations of this exploratory pilot study should be noted. First, as a naturalistic pilot study, the duration and timing of dog–patient contact were not experimentally manipulated or standardized; events occurred spontaneously based on clinical flow. Second, while skin conductance synchrony provides a sensitive measure of sympathetic coordination, physiological synchrony is context-dependent and does not inherently distinguish between positive affiliative attunement and shared co-arousal or stress (Gernert et al., 2024). Future research should incorporate continuous video coding or post-session self-reports (e.g., therapeutic alliance or affect measures) to map physiological synchrony onto specific behavioral states and subjective emotional experiences.

Additionally, future studies should examine triadic physiological synchrony by simultaneously recording autonomic signals from all three participants: patient, therapist, and therapy dog (Koskela et al., 2024). Tracking dog–human physiological coupling alongside therapist– patient synchrony would provide a more complete understanding of how autonomic states cascade across species during canine-assisted psychotherapy (Suttkus et al., 2026; Vierimaa and Pyyhtinen, n.d.).

### Conclusion

This study provides initial evidence that direct physical contact between a patient and a therapy dog during canine-assisted psychotherapy is followed by significant increases in both individual autonomic arousal and therapist–patient physiological synchrony. By using an event-based approach with pseudo-event controls, we demonstrate that contact with a therapy dog acts as a specific interactional event that promotes temporal coordination between therapist and patient. These findings support the view that therapy dogs serve as active social catalysts within the therapeutic triad, creating physiological conditions that may facilitate interpersonal attunement and strengthen the therapeutic relationship.

